# *Chlamydomonas* γ-tubulin mutations reveal a critical role for γ-TuRC in maintaining the stability of centriolar microtubules

**DOI:** 10.1101/2025.03.18.643570

**Authors:** Yuki Nakazawa, Naoto Kubota, Mao Horii, Akira Noga, Yoshikazu Koike, Hiroko Kawai-Toyooka, Hideo Dohra, Katsushi Yamaguchi, Shuji Shigenobu, Ken-ichi Wakabayashi, Masafumi Hirono

## Abstract

The centriolar triplet microtubule consists of an A-tubule with 13 protofilaments, and B-and C-tubules, each with 10 protofilaments. Although the formation of the triplets has been shown to require γ-tubulin, its specific role in the formation of each tubule remains elusive. We isolated two novel *Chlamydomonas reinhardtii* mutants, *bld13-1* and *bld13-2*, each expressing γ-tubulin with a single amino-acid substitution (T292I or E89D). Similar to known centriole-deficient mutants, both mutants exhibited defects in ciliary assembly, nuclear number, as well as the number and orientation of cytoplasmic microtubules. Genetic analyses of the mutants, along with the expression of the mutant γ-tubulins in the wild-type cells, suggested that both mutants exert dominant-negative effects over wild-type γ-tubulin. Interestingly, although the centrioles in these mutants retained the typical nine triplet structure, their triplets frequently lacked several protofilaments in specific regions of the A- and C-tubules. The protofilament loss occurs more frequently in the proximal region of the centriole. These structural defects suggest a critical role for γ-tubulin in maintaining the stability of the A- and C-tubules of centriolar triplets.

**Summary statement:** Novel *Chlamydomonas* γ-tubulin mutations cause a partial loss of protofilaments in centriolar microtubules, indicating a critical role of γ-tubulin in structural stabilization of triplet microtubules.

## Introduction

Triplet microtubules are a specialized type of microtubule found exclusively in centrioles, structures with 9-fold symmetry that serve as the core of the centrosome and the basal bodies of cilia (flagella). Each triplet consists of an A-tubule, composed of 13 protofilaments arranged in a circle, and B- and C-tubules, each composed of 10 protofilaments arranged in an arc. When centrioles function as basal bodies, the outer doublet microtubules of the axoneme extend from the A- and B-tubules (Wang and Stearns, 2017; Osinka et al., 2019; Le Guennec et al., 2021). The unique structure of the centriolar triplet microtubules is highly conserved across eukaryotic organisms, yet its assembly mechanism remains poorly understood (Azimzadeh and Marshall, 2010; Wang and Stearns, 2017; Guichard et al., 2023).

Similar to the assembly of cytoplasmic microtubules, the triplet assembly likely involves the γ-tubulin ring complex (γ-TuRC) (Guichard et al., 2010; Schweizer and Lüders, 2021; Mukhopadhyay et al., 2026), a major nucleator of tubulin polymerization (Oakley et al., 2015; Wu and Akhmanova, 2017). It is composed of 14 γ-tubulin molecules arranged in an incomplete ring, along with GCP2∼6, MZT1, and other proteins. This structure suggests that it serves as a template for microtubule nucleation, positioning each γ-tubulin at the minus end of a protofilament during nucleation (Liu et al., 2021; Thawani and Petry, 2021; Zupa et al., 2021; Sulimenko et al., 2022). In animal cells, numerous γ-TuRCs within the pericentriolar materials (PCM) of the centrosome nucleate cytoplasmic microtubules, enabling the centrosome to function as the microtubule-organizing center (MTOC) (Oakley et al., 2015; Wu and Akhmanova, 2017).

In fact, the γ-tubulin or γ-TuRC has been reported to be essential for centriole formation in many organisms (Ruiz et al., 1999; Shang et al., 2002; Dammermann et al., 2004; Gogendeau et al., 2011; Bahtz et al., 2012; Cota et al., 2017). By analogy to its role in cytoplasmic microtubule assembly, the γ-TuRC is thought to function in the nucleation of centriolar triplet microtubules as well. Supporting this hypothesis, cryo-electron microscopy of isolated human centrosomes revealed a conical structure, likely a γ-TuRC, capping the proximal end of the A-tubule in the immature centriole (Guichard et al., 2010). However, the γ-TuRC’s nucleation activity for centriolar microtubules has not been directly demonstrated, and its contribution to triplet microtubule assembly and/or maintenance, particularly of the B- and C-tubules, remains unclear.

Besides its prominent localization to the PCM, γ-TuRC is also found in various centriolar sites: in the proximal region of the triplet microtubule wall, on the outer and inner (luminal) surfaces of the microtubule wall, and on the distal appendages (Silflow et al., 1999; Schweizer and Lüders, 2021; Kalbfuss and Gönczy, 2023; Laporte et al., 2024). While the specific function of γ-TuRC at each site is unclear, a recent study in human cells showed that the depletion of proteins that localize γ-TuRCs to the inner surfaces of the centriole microtubules destabilizes the centrioles and impairs ciliary assembly (Schweizer et al., 2021). This suggests that the γ-TuRCs within the centriole lumen play a crucial role in maintaining centriole structural integrity.

In this study, we isolated two novel *Chlamydomonas reinhardtii* strains carrying missense mutations in the single copy γ-tubulin gene, which appear to exert dominant-negative effects. These are the first viable mutants of the γ-tubulin gene identified in *Chlamydomonas reinhardtii*. Similar to previously reported centriole-deficient mutants, these mutants exhibited defects in ciliary assembly. Intriguingly, their centrioles frequently lacked some of the protofilaments in the A- and C-tubules of triplet microtubules. These phenotypes and the predicted positions of the mutations in the γ-tubulin molecule support the idea that γ-tubulin functions as a γ-TuRC component to stabilize triplet microtubules.

## Results

### Novel γ-tubulin mutants of *Chlamydomonas*

We isolated two independent *Chlamydomonas* strains that showed defects in ciliary assembly and nuclear segregation, which are phenotypes characteristic of known centriole-deficient mutants (Fig. 1 A, B; Table 1) (Ehler et al., 1995; Dutcher and Trabuco, 1998; Matsuura et al., 2004; Hiraki et al., 2007; Nakazawa et al., 2007). As genetic analyses mapped these mutations to the same locus on Linkage Group VI (Chromosome 6), we named the mutants *bld13-1* and *bld13-2* (Fig. 1C). Further linkage analyses showed that the causal mutations exhibited no recombination with a genetic marker designed within the γ-tubulin gene; among 188 progenies examined, no recombinants were detected. Whole genome sequencing of *bld13-1*, and PCR-based sequencing of the *bld13-2* γ-tubulin gene identified point mutations that cause single amino-acid substitutions (*bld13-1*, T292I; *bld13-2*, E89D) in the γ-tubulin gene, a single-copy gene in *Chlamydomonas* (Fig. 1D). The original amino acids at these mutation sites are widely conserved among organisms including single-cell organisms and humans (Fig. 1D). To infer possible effects of these mutations on the γ-tubulin structure, we compared AlphaFold2-predicted models of the *bld13-1* and *bld13-2* γ-tubulins with that of the wild type (Fig. S1A) (Jumper et al., 2021). The models suggest no predominant changes in the molecular conformation, except for a slight change in a small region on the molecular surface of *bld13-1* γ-tubulin.

**Fig. 1.**
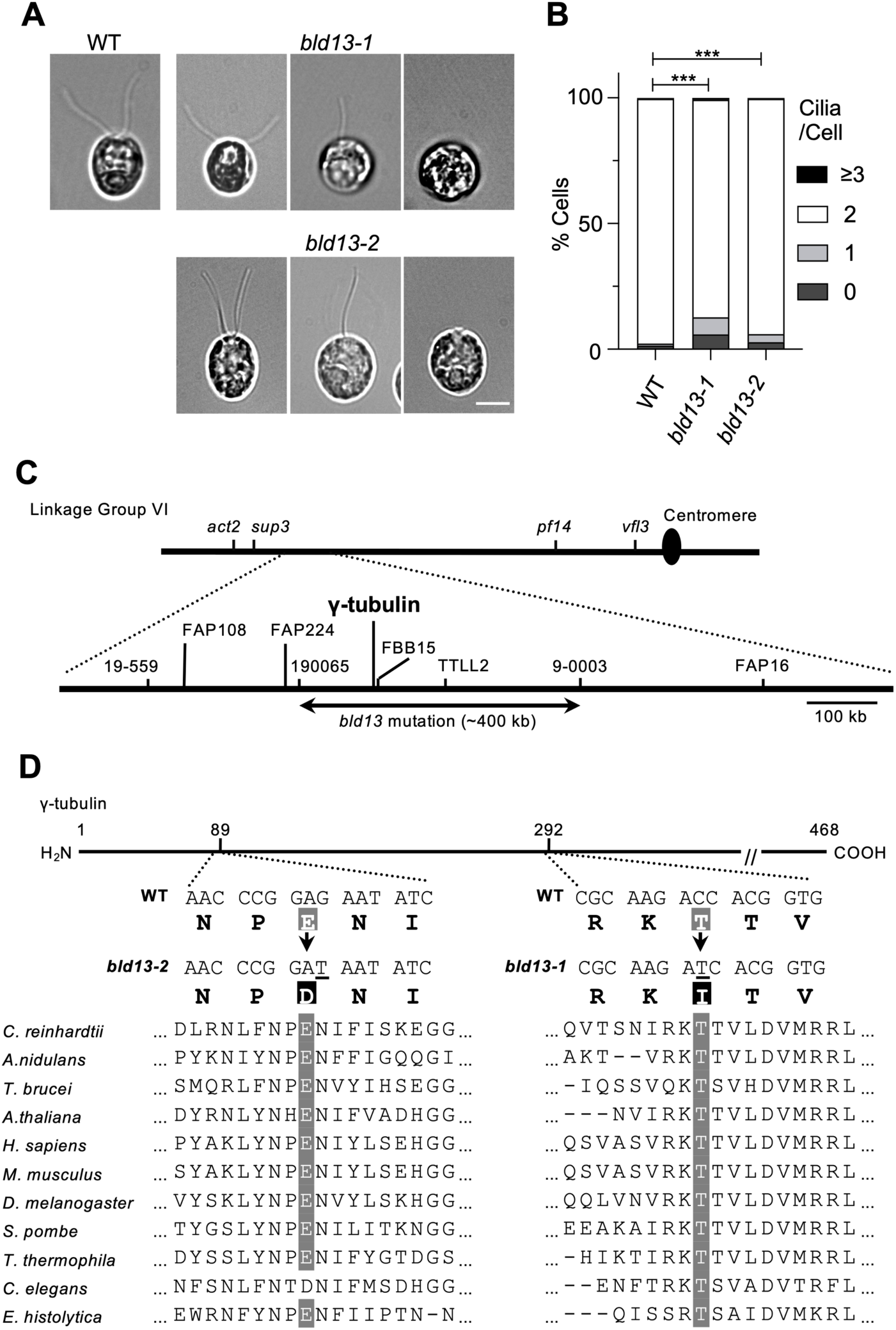
Novel *Chlamydomonas* mutants *bld13-1* and *bld13-2*. (A) Phase contrast images of wild-type, *bld13-1* and *bld13-2* cells. Both *bld13* mutants contain cells with no or one cilium in addition to cells with two cilia. Scale bar: 5 µm. (B) Percentages of cells with zero, one, or two cilia in wild type (WT), *bld13-1*, and *bld13-2*. Statistical analysis was performed using Fisher’s exact test (***P<0.001; n>1500). (C) A genetic map showing a part of Linkage Group VI. The *bld13-1* and *bld13-2* mutations were mapped to an ∼400-kb region in this linkage group by AFLP-based analyses. The upper line shows the corresponding locus on the genetic map constructed by Kathir et al. (Kathir et al., 2003). The lower line shows positions of genes around this region (JGI Phytozome v 13, *Chlamydomonas* database v 5.6 https://phytozome-next.jgi.doe.gov/). Numbers 19-559, 190065, and 9-0003 represent primer sets used for PCR. FAP16, FAP108, and FAP224 are genes of flagellar associated proteins (Pazour et al., 2005), FBB15 is a flagella/basal body protein gene (Li et al., 2004), and TTLL2 is a putative tubulin tyrosine ligase gene (Cre06.g300250, JGI Phytozome v 13, *Chlamydomonas* database v 5.6, https://phytozome-next.jgi.doe.gov/). (D) The positions of the *bld13-1* and *bld13-2* mutations in the γ-tubulin amino acid sequence of *Chlamydomonas* and corresponding sequences in other organisms. GenBank accession numbers are as follows: *C. reinhardtii*, AAA82610.1; *A. nidulans*, CAA33507.1; *T. brucei*, XP_843687.1; *A. thaliana*, NP_191724.1; *H. sapiens*, NP_001061.2; *M. musculus*, NP_598785.1; *D. melanogaster*, NP_476804.1; *S. pombe*, NP_596147.1; *T. thermophila*, AAB65830.1; *C. elegans*, NP_499131.1; *E. histolytica*, AAC08441.1.

**Table 1.** Abnormal nuclear segregation in the *bld13* mutants.

| Strain | Number of nuclei/cell (n>2200) |  |  |  |
| --- | --- | --- | --- | --- |
|  | 0 | 1 | 2 or 3 |  |
| wild type | 0.61 | 99.30 | 0.09 |  |
| *bld12-1* | 3.70 | 90.20 | 6.11 |  |
| *bld13-1* | 1.24 | 95.34 | 3.43 |  |
| *bld13-2* | 0.64 | 98.39 | 0.97 | (%) |

To computationally assess their functional impact, AlphaMissense was applied to human γ-tubulin (UniProt ID P23258) (Cheng et al., 2023). The equivalent E89D and T288I (corresponding to *Chlamydomonas* E89D and T292I, respectively) substitutions were both classified as ‘likely pathogenic’ (scores: 0.9508 and 0.9956, respectively).

### The *bld13-1* and *bld13-2* mutations are consistent with a dominant-negative mechanism

The defective phenotypes of the *bld13* mutants were not rescued by expression of a hemagglutinin (HA)-tagged γ-tubulin gene (γ-tub(WT)-HA). In addition, expression in wild-type cells of an HA-tagged γ-tubulin gene carrying either the *bld13-1* or *bld13-2* mutation (WT::γ-tub(T292I)-HA or WT::γ-tub(E89D)-HA), driven by the endogenous promoter, caused defects in ciliary assembly similar to those observed in the mutants (Fig. 2 A, B). Together, these results suggest that the mutant γ-tubulins may interfere with wild-type γ-tubulin function, consistent with a dominant-negative effect. To test this possibility under equal gene dosage of wild-type and mutant alleles, we produced diploid strains by crossing the wild type with *bld13-1* or *bld13-2* (WT/*bld13-1*, WT/*bld13-2*), and counted the percentage of cells with no or only one cilium in the two heterozygous strains as well as in the homozygous diploid strain (*bld13-1*/*bld13-2*). The percentages were significantly higher in the heterozygous or homozygous diploid strains carrying the *bld13* mutation than in the wild-type diploid strain (WT/WT) (Fig. 2C). These results support the interpretation that the *bld13-1* and *bld13-2* mutations interfere with wild-type γ-tubulin function in a dominant-negative manner.

**Fig. 2.**
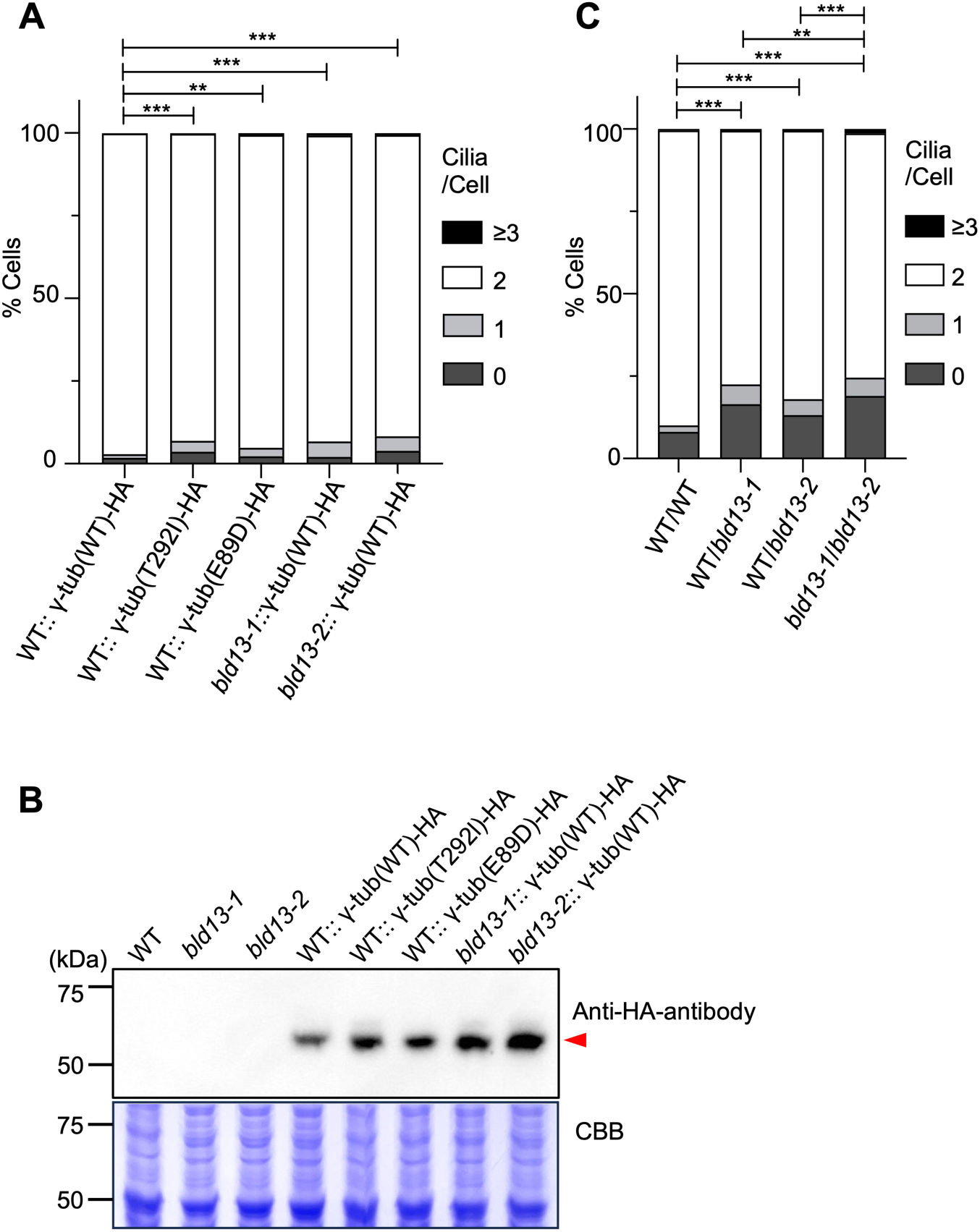
Evidence supporting dominant-negative effects of the *bld13* mutations on ciliary assembly. (A) Percentages of cells with zero, one, or two cilia in strains ectopically expressing HA-tagged γ-tubulin from genomic fragments under the control of the endogenous γ-tubulin promoter. These strains include a wild type strain expressing an HA-tagged γ-tubulin (WT::γ-tub(WT)-HA), a wild type strain expressing the *bld13-1* γ-tubulin (WT::γ-tub(T292I)-HA), a wild type strain expressing the *bld13-2* γ-tubulin (WT::γ-tub(E89D)-HA), a *bld13-1* strain expressing the HA-tagged γ-tubulin (*bld13-1*::γ-tub(WT)-HA), and a *bld13-2* strain expressing the HA-tagged γ-tubulin (*bld13-2*::γ-tub(WT)-HA). Statistical analysis was performed using Fisher’s exact test (***P < 0.001 for WT::γ-tub(WT)-HA vs *bld13-1*::γ-tub(WT)-HA, *bld13-2*::γ-tub(WT)-HA, and WT::γ-tub(T292I)-HA; **P = 0.0013 for WT::γ-tub(WT)-HA vs WT::γ-tub(E89D)-HA; n>1500). (B) Western blot analyses of whole cell lysates (10 μg/lane) of the wild type (WT), *bld13-1*, *bld13-2*, WT::γ-tub(WT)-HA, WT::γ-tub(T292I)-HA, WT::γ-tub(E89D)-HA, *bld13-1*::γ-tub(WT)-HA, and *bld13-2*::γ-tub(WT)-HA cells. The upper panel shows immunoblotting with an anti-HA antibody detecting γ-tubulin(WT)-HA (55 kDa; arrowhead), and the lower panel shows Coomassie Brilliant Blue (CBB) staining as a loading control. (C) Percentages of cells with zero, one, or two cilia in diploid strains of the wild type (WT/WT), heterozygous *bld13-1* (WT/*bld13-1*), heterozygous *bld13-2* (WT/*bld13-2*), and *bld13-1*/*bld13-2*. Statistical analysis was performed using Fisher’s exact test (***P < 0.001 for WT/WT vs each mutant genotype and for WT/*bld13-2* vs *bld13-1*/*bld13-2*; **P = 0.0036 for WT/*bld13-1* vs *bld13-1*/*bld13-2*; n >1500).

### γ-Tubulin localizes to centrioles and their surrounding regions independently of *bld13* mutations

In the interphase *Chlamydomonas* cell, two mature centrioles (basal bodies) and two immature centrioles (probasal bodies) are present at the base of the two cilia, forming an organizing center for four stable rootlet microtubule bundles and non-rootlet cytoplasmic microtubules (Fig. 3A; Holmes and Dutcher, 1989). A positional marker for centrioles, SAS-6 localizes to the proximal ends of both mature and immature centrioles and is therefore observed as four discrete fluorescent spots.

**Fig. 3.**
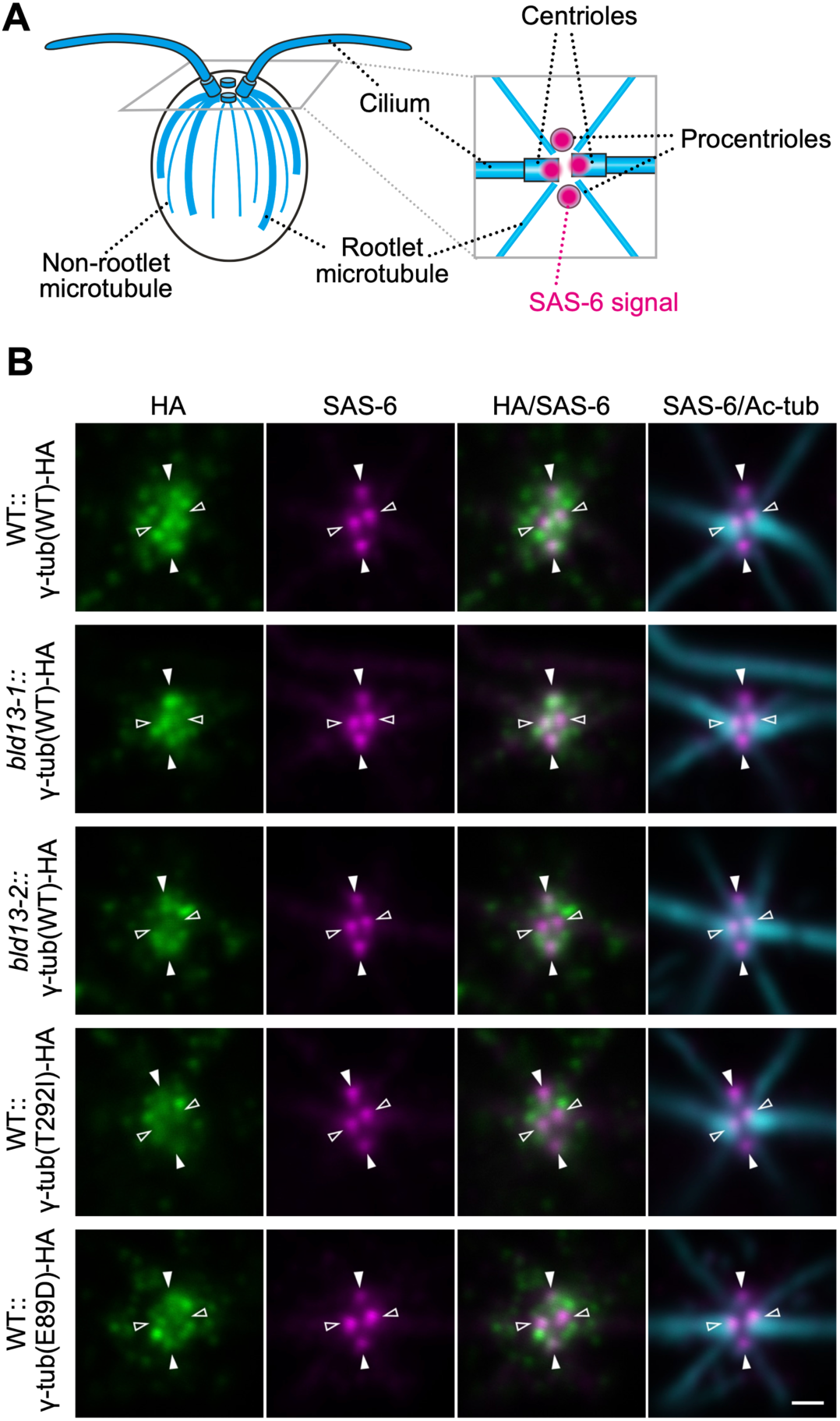
Localization of γ-tubulin in the presence or absence of *bld13* mutations. (A) Schematic diagram showing microtubule organization and centriole positioning in the *Chlamydomonas* cell. (B) Immunofluorescence images of WT::γ-tub(WT)-HA, *bld13-1*::γ-tub(WT)-HA, *bld13-2*::γ-tub(WT)-HA, WT::γ-tub(T292I)-HA, and WT::γ-tub(E89D)-HA cells stained with anti-HA (HA, green), anti-SAS-6 (SAS-6, magenta), and anti-acetylated tubulin (Ac-tub, cyan) antibodies. Merged images show HA and SAS-6 signals (HA/SAS-6) or SAS-6 and acetylated tubulin signals (SAS-6/Ac-tub). Scale bar: 500 nm. Filled arrowheads indicate procentrioles, and open arrowheads indicate mature centrioles. Representative images are shown; similar localization patterns were consistently observed.

In wild-type and *bld13* cells expressing HA-tagged wild-type γ-tubulin (Fig. 3B; WT::γ-tub(WT)-HA, *bld13-1*::γ-tub(WT)-HA, and *bld13-2*::γ-tub(WT)-HA), HA signals were detected in regions adjacent to the four centrioles, which are marked by SAS-6 fluorescence. Most HA signals were observed as punctate fluorescence in the pericentriolar region, together with diffuse fluorescence distributed throughout this region. Some SAS-6 fluorescent spots overlapped with punctate HA signals, whereas the remaining SAS-6 spots overlapped with the diffuse HA fluorescence.

In wild-type cells expressing HA-tagged mutant γ-tubulin (Fig. 3B; WT::γ-tub(T292I)-HA and WT::γ-tub(E89D)-HA), the overall HA localization patterns were essentially indistinguishable from those observed in the cells expressing HA-tagged wild-type γ-tubulin. In all cases, HA signals were detected in association with SAS-6–identified centrioles, together with prominent punctate or diffuse signals in the surrounding regions.

These observations indicate that γ-tubulin–HA signals, although not always strongly concentrated, overlap with both mature and immature centrioles, particularly in their pericentriolar regions. Apparently, the *bld13-1* and *bld13-2* mutations do not grossly alter the subcellular localization pattern of HA-tagged γ-tubulin.

### Cytoplasmic microtubule organization is aberrant in *bld13* cells

In mammalian cells, γ-tubulin localizes to the pericentriolar material (PCM) of the centrosome and functions in the nucleation and organization of cytoplasmic microtubules (Oakley et al., 2015; Wu and Akhmanova, 2017). In *Chlamydomonas* cells, although a distinct PCM has not been observed, functional analogs of PCMs are most likely attached to the centrioles and serve as the site where γ-tubulin functions, as rootlet and non-rootlet cytoplasmic microtubules extend from the proximity of the centrioles (Fig. 3A, Holmes and Dutcher, 1989). To evaluate the effects of the *bld13* mutations on the intracellular microtubule-organization, we performed immunofluorescence microscopy using two kinds of tubulin antibodies. One was an anti–acetylated tubulin antibody, which primarily recognizes axonemal and rootlet microtubules (Fig. 4A), and the other was an anti–α-tubulin antibody, which recognizes both rootlet microtubules and non-rootlet cytoplasmic microtubules (Fig. 4B).

**Fig. 4.**
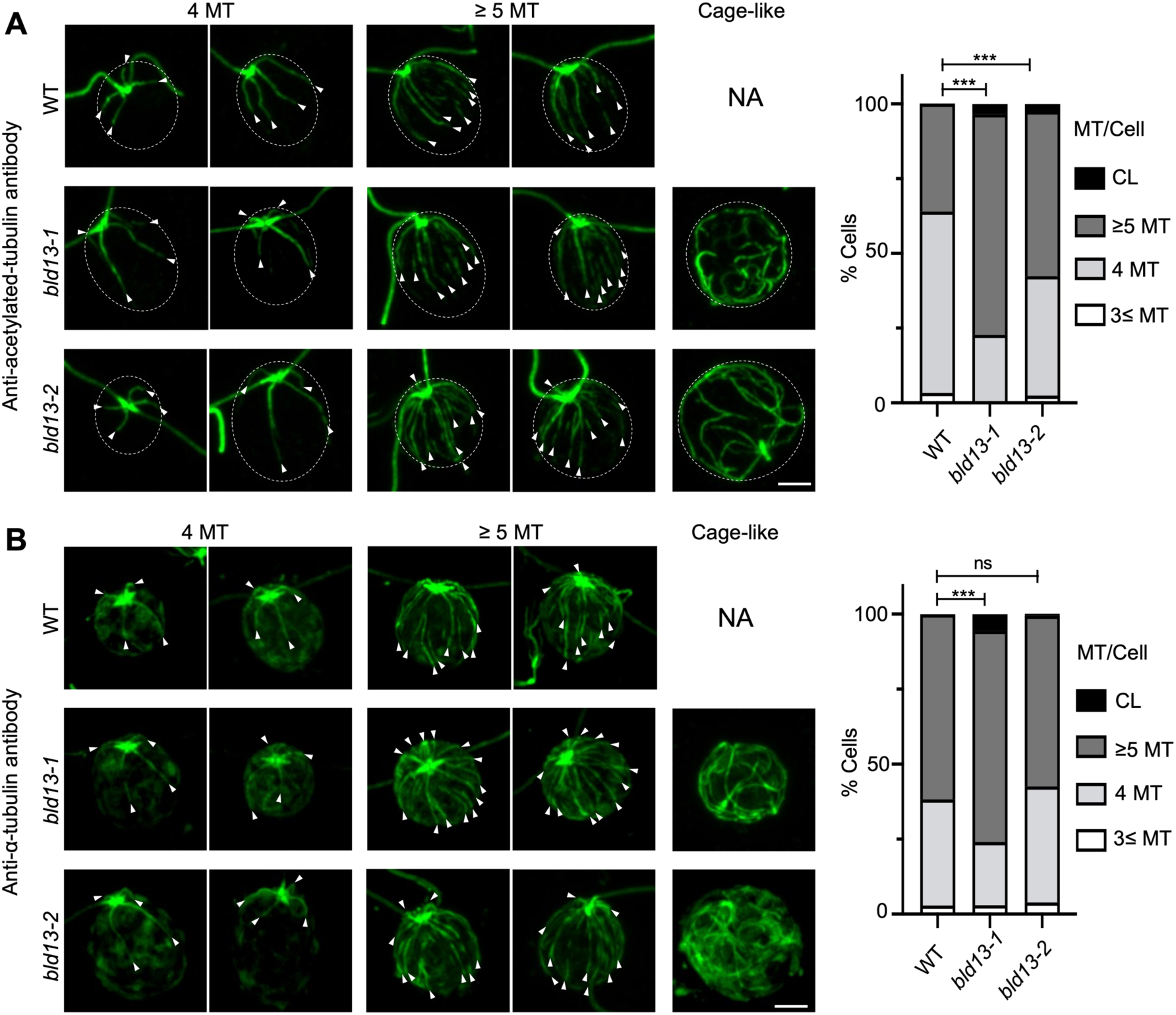
Effects of *bld13* mutations on microtubule organization in *Chlamydomonas* cells. All images are shown as Z-projection images of confocal optical sections. (A) Immunofluorescence images of wild-type, *bld13-1*, and *bld13-2* cells stained with anti-acetylated α-tubulin antibody, which recognizes rootlet, axonemal, and centriolar microtubules, but not non-rootlet cytoplasmic microtubules (Holmes and Dutcher, 1989). Cells were classified into four categories based on acetylated microtubule organization: cells with three or fewer acetylated microtubules (≤3 MT), cells with four rootlets (4 MT), cells with five or more acetylated microtubules (≥5 MT), and cells lacking an obvious microtubule-organizing center but exhibiting a cage-like acetylated microtubule (cage-like, CL). Representative images of each category are shown. Arrowheads indicate rootlet or acetylated microtubules with unclear rootlet identity. The graph shows the frequency of each category. (B) Immunofluorescence images of wild-type, *bld13-1*, and *bld13-2* cells stained with an anti–α-tubulin antibody, which labels non-rootlet cytoplasmic microtubules in addition to rootlet, axonemal, and centriolar microtubules. Cells were classified using the same criteria as in (A), and the frequency of each category is shown in the graph. Statistical analysis was performed using Fisher’s exact test (***P < 0.001; ns, not significant; n>120). Scale bars: 2 µm.

When cells were stained with the anti–acetylated α-tubulin antibody, ∼60% of wild-type cells exhibited a typical cruciform rootlet arrangement of four microtubule bundles, consistent with the canonical rootlet organization (Fig. 4A). In the remaining ∼40% of wild-type cells, five or more acetylated microtubule structures were observed, but it was not possible to definitely identify these microtubules as rootlets. In contrast, the proportion of cells with five or more acetylated microtubules were significantly increased in the mutants, reaching 75% in *bld13-1* and 55% in *bld13-2*. Notably, a subset of mutant cells displayed cage-like cytoplasmic microtubules that did not originate from an obvious organizing center; such cells were observed at frequencies of 3.4% in *bld13-1* and 2.4% in *bld13-2*, but were never observed in the wild type.

Staining with the anti–α-tubulin antibody allowed visualization of non-rootlet cytoplasmic microtubules in addition to rootlet microtubules (Fig. 4B). With this antibody, even wild-type cells frequently exhibited five or more microtubules, making the distinction between wild-type and mutant cells less pronounced than with anti–acetylated α-tubulin staining. However, in both *bld13* mutant strains, but not in wild type, cells exhibiting cage-like microtubule arrays were again detected. In these cells, neither acetylated nor non-acetylated microtubules appeared to originate from a discrete microtubule-organizing center.

Taken together, these observations indicate that the *bld13-1* and *bld13-2* γ-tubulins are likely to have some defects in organizing microtubules in *Chlamydomonas* cell bodies.

### Centriolar triplet microtubules frequently lack protofilaments in *bld13*

The phenotypes of *bld13* mutants, namely, defective in ciliary assembly, nuclear segregation, and rootlet microtubule organization, are shared by various *Chlamydomonas* mutants having structural defects in the centriole (Ehler et al., 1995; Dutcher and Trabuco, 1998; Matsuura et al., 2004; Hiraki et al., 2007; Nakazawa et al., 2007). Therefore, we examined the centriole structure in *bld13* cells by thin-section electron microscopy. Longitudinal images of *bld13-1* and *bld13-2* centrioles appeared normal (Fig. 5A), although some of those centrioles did not attach axonemes and therefore apparently failed to nucleate ciliary growth. Strikingly, however, obvious defects were observed in cross-sectional images of centrioles in both mutants: some triplet microtubules partially lacked protofilaments (Fig. 5B). In most cases, two to seven adjacent protofilaments were lost from particular positions in the A- and C-tubules, but not in the B-tubules. These abnormalities were never observed in wild-type centrioles (Table 2). To evaluate the cross-sectional positions in the *bld13* centrioles where protofilament loss frequently occurs, the cross-sectional images of the A- or C-tubules were divided into 6 or 12 sectors, and the occurrence of missing protofilaments was scored in each sector. The results showed that the protofilament loss frequently occurred in the sectors that contain protofilaments No. A1-A6 of the A-tubule and No. C1-C7 of the C-tubule in both *bld13-1* and *bld13-2* (the numbering is after Linck and Stephens, 2007 and Li et al., 2019). These data indicate that protofilament loss occurred non-randomly, with a significantly biased distribution in both A- and C-tubules (Fig. 5C).

**Fig. 5.**
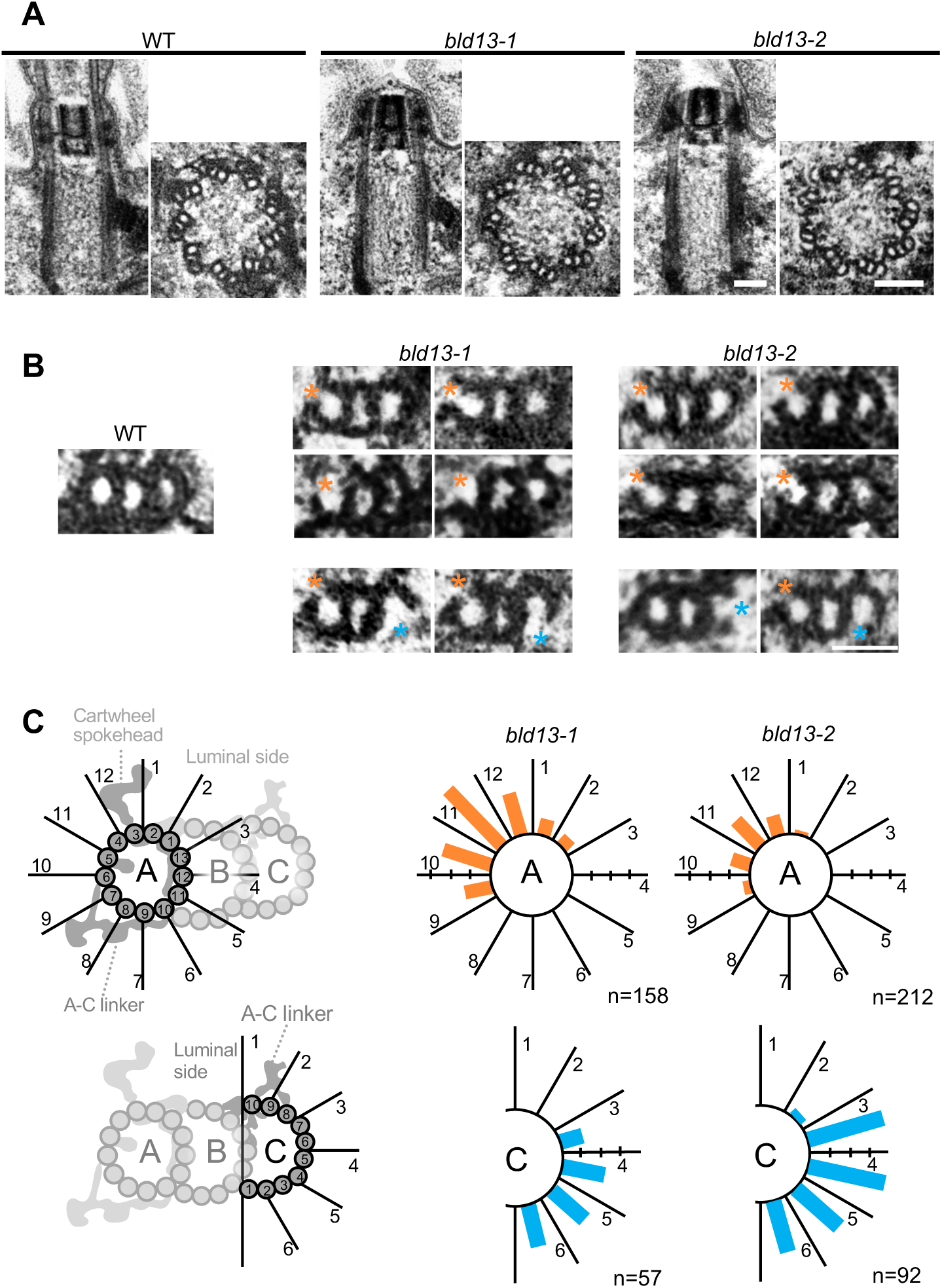
Partial loss of protofilaments in *bld13* centriolar triplet microtubules. (A) Electron microscopic images of longitudinal and cross sections of centrioles in WT, *bld13-1*, and *bld13-2* cells. Scale bars: 100 nm. (B) Magnified views of cross-sections of the triplet microtubules in WT, *bld13-1*, and *bld13-2* centrioles. Asterisks indicate the positions of protofilament loss. Scale bar: 40 nm. (C) Distributions of the protofilament loss in A- and C-tubules of *bld13-1* and *bld13-2* triplet microtubules. A cross-sectional image of the A-or C-tubule was divided into 12 or 6 sectors as shown in schematic diagrams (left, see Materials and Methods). MIPs (in the lumen of the A-tubule), the A-C linker, and the cartwheel spokehead, are drawn in grey (Li et al., 2019). The length of each bar in the polar histograms (right) represents the relative frequency of protofilament loss at the corresponding sector, expressed as a percentage of the total number of observed triplet microtubule images. The x-axis of each graph is shown in 5% increments. Statistical analysis was performed using the Hodges–Ajne test, showing that all distributions of the protofilament loss are significantly non-uniform: *bld13-1* A-tubule (P < 0.0001; median: sector 11), *bld13-2* A-tubule (P < 0.0001; median: sector 11), *bld13-1* C-tubule (P < 0.0001; median: sector 4), *bld13-2* C-tubule (P < 0.0001; median: sector 4).

**Table 2.**
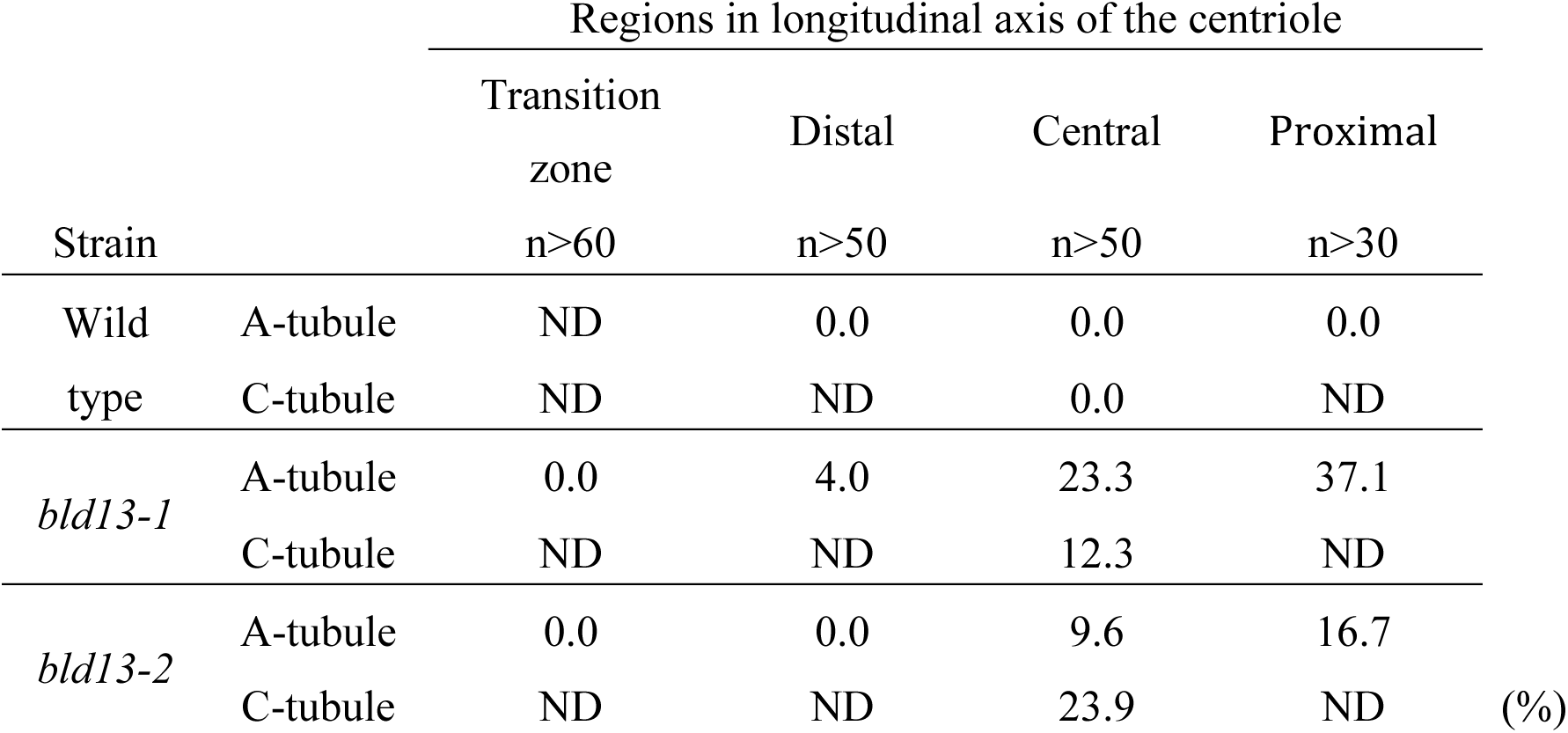
Frequencies of triplet images with partial protofilament loss (%). Percentages indicate the proportion of observed triplet microtubule images showing loss of one or more protofilaments.

| Strain |  | Regions in longitudinal axis of the centriole |  |  |  |  |
| --- | --- | --- | --- | --- | --- | --- |
|  |  | Transition zone | Distal | Central | Proximal |  |
|  |  | n>60 | n>50 | n>50 | n>30 |  |
| Wild type | A-tubule | ND | 0.0 | 0.0 | 0.0 |  |
|  | C-tubule | ND | ND | 0.0 | ND |  |
| *bld13-1* | A-tubule | 0.0 | 4.0 | 23.3 | 37.1 |  |
|  | C-tubule | ND | ND | 12.3 | ND |  |
| *bld13-2* | A-tubule | 0.0 | 0.0 | 9.6 | 16.7 |  |
|  | C-tubule | ND | ND | 23.9 | ND | (%) |

The frequency of the protofilament loss in the A-tubule seemingly varied between the proximal and the distal regions of the centriole. To objectively show this difference, we counted the number of missing protofilaments in four groups of axonemal images that most likely represented distinct positions of the centriole: 1) (transition zone) images with the characteristic star-like structure in the lumen; 2) (distal region) images with distal appendages; 3) (proximal region) images with cartwheels; and 4) (central region), images other than the above three groups (Table 2) (Geimer and Melkonian, 2004; Le Guennec et al., 2021). The results showed that the protofilament loss of the A-tubule was more frequently observed in the proximal region than in the central or distal regions. No protofilament loss was observed in the transition zone. We were unable to examine the regional difference of protofilament loss in the C-tubule, because its image was obscured by electron-dense material near the proximal region and was often missing in the distal region.

## Discussion

In this study we demonstrated that two novel mutations in the γ-tubulin gene in *Chlamydomonas* cause partial loss of protofilaments in centriolar triplet microtubules while having relatively mild effects on the overall centriole structure. The ultrastructural defects observed in the mutant centrioles highlight the crucial role of γ-tubulin in maintaining the stability of triplet microtubules. This study provides the first isolation and characterization of γ-tubulin mutants in *Chlamydomonas*.

### Effects of *bld13* mutations on the γ-TuRC assembly

As γ-tubulin is an essential component of the microtubule-organizing center (MTOC), required for fundamental cellular processes like cell division (Paz and Lüders, 2018; Sulimenko et al., 2022), loss of γ-tubulin function results in lethality in many model organisms (Sunkel et al., 1995; Hendrickson et al., 2001; Yuba-kubo et al., 2005; Pastuglia et al. 2006). It is fortunate that the *bld13* mutants reported in this study have only mild phenotypic aberrations and thus can provide a tractable system to dissect the *in vivo* functions of γ-tubulin.

In many eukaryotes, γ-tubulin constitutes two kinds of complexes, one termed the γ-tubulin small complex (γ-TuSCs) and the other termed the γ-tubulin ring complex (γ-TuRC). The γ-TuSC is formed by association between γ-tubulin and two subunits, GCP2 and GCP3; the γ-TuRCs is formed by oligomerization of γ-TuSCs and association with GCP4∼6 and MZT1 (Tovey and Conduit, 2018; Böhler et al., 2021). Because *Chlamydomonas* has genes coding for homologs of GCP2∼4 and MZT1, γ-tubulin of this organism most likely also functions in microtubule nucleation by forming γ-TuRCs.

Both *bld13* mutations in the γ-tubulin gene cause single amino-acid substitutions that do not change side-chain charge (Fig. 1D). Consistently, AlphaFold2 predicts no significant changes in the overall molecular structure (Fig. S1A). The predicted minor structural effects of these mutations may be correlated with the mild phenotypic abnormalities in the mutants: the mutant γ-tubulins localize to both centrioles and the surrounding pericentriolar region as in wild-type cells (Silflow et al., 1999), and cytoplasmic microtubules and ciliary axonemes are formed in most of the mutant cells (Fig. 1-3). On closer inspection, however, the *bld13* mutations were associated with three detectable abnormalities in microtubule structures in a subset of cells. One abnormality was an increased population of cells exhibiting elevated numbers of cytoplasmic microtubules, most clearly detected by staining for acetylated α-tubulin (Fig. 4). A second abnormality, observed in a small fraction of cells, was the appearance of cytoplasmic microtubules that lacked an obvious microtubule-organizing center (Fig. 4). A third abnormality was a partial loss of protofilaments in centriolar microtubules (Fig. 5).

We speculate that these defects could result from some instability in the association of the γ-tubulin molecules within the γ-TuRC. This is because the mutated amino acid residues in both *bld13* mutants are located on the molecular surface at the region where adjacent γ-tubulin molecules are presumed to interact within the complex (Fig. S1B and C; Inclán and Nogales, 2000; Aldaz et al., 2005; Wieczorek et al., 2020). In addition, AlphaMissense predicts that the equivalent substitutions in human γ-tubulin are “likely pathogenic” (Cheng et al., 2023), further supporting the functional importance of these residues. We propose that γ-tubulin molecules are stably held in the complex by inter-subunit interaction at the contact sites, and the *bld13* mutations partially disrupt this interaction.

Diploid analyses further support the dominant-negative mode of the *bld13* mutations, consistent with the results of the γ-tubulin expression experiments (Fig. 2). The presence of both wild-type and mutant γ-tubulin in heterozygous diploids suggest that mutant subunits are incorporated into the γ-TuRC together with wild-type subunits, thereby reducing the overall functional integrity of the complex. Moreover, the slightly severer phenotype observed in the *bld13-1*/*bld13-2* homozygous diploid suggests that the coexistence of the two mutations impairs inter-subunit contact surfaces more strongly within the γ-TuRC. We speculate that weakened contact between γ-tubulins may compromise γ-TuRC stability, leading to the defective cellular microtubule organization and the partial loss of protofilaments observed in *bld13* mutants.

It remains unclear how the structural defects in the *bld13* γ-TuRC results in the abnormal organization of the rootlet and non-rootlet microtubules (Fig. 4). Since other centriole-deficient mutants we studied previously, *bld10* and *bld12*, also exhibited abnormal rootlet microtubule arrangement (Matsuura et al., 2004; Nakazawa et al., 2007), it is possible that the microtubule organization in *Chlamydomonas* is extremely susceptible to the structural abnormality of the centriole.

### γ-tubulin functions in stabilization of centriolar microtubules

The triplet microtubules of *bld13* centrioles frequently lack part of the protofilaments (Fig. 5). This type of microtubule defects is unexpected because the lateral interaction between protofilaments is likely essential for the stability of microtubules, and loss of a protofilament would lead to depolymerization of whole microtubules (Gudimchuk and McIntosh, 2021). However, the triplet microtubules in *bld13*, even when lacking protofilaments, do not fully depolymerize. We speculate that these microtubules are stabilized by some microtubule-associated structures or proteins, such as the A–C linkers, microtubule inner proteins (MIPs), and the inner-scaffolds of the centriole (Li et al., 2019; Le Guennec et al., 2020). It is interesting to note that the protofilament loss is frequently observed in the region centered on A4 and the region C1-C7, where protofilaments are apparently not associated with those structures or MIPs (Fig. 5C).

Protofilament loss in the A-tubule was observed more frequently in the proximal region than in the central or distal region in both *bld13-1* and *bld13-2* centrioles (Table 2), suggesting that the loss was caused by depolymerization from the proximal end of previously formed protofilaments. *In vitro* studies have shown that the γ-TuRC caps the microtubule minus end for a short period and suppresses both polymerization and depolymerization (Wiese and Zheng, 2000; Berman et al., 2023). In centriolar triplet microtubules *in vivo*, their minus ends may be mostly capped by the γ-TuRC, since γ-tubulin has been localized near the proximal ends of the mature centrioles in *Tetrahymena*, *Paramecium*, and mammals (Fuller et al., 1995; Klotz et al., 2003; Joachimiak et al., 2018). Moreover, cryo-electron tomography of human procentrioles has shown that the A-tubule proximal ends have a conical shape resembling the γ-TuRC-capped microtubule end observed *in vitro* (Guichard et al., 2010). Thus, our observations support the possibility that the γ-TuRC caps the proximal end of the A-tubule to suppress depolymerization, and that this capping function may be compromised in the *bld13* centrioles due to weakened interaction between γ-tubulin molecules.

Electron microscopy of the *bld13* centrioles revealed partial loss of protofilaments also in the C-tubules (Fig. 5 B,C). The mechanism underlying formation and maintenance of B- and C-tubules in centrioles remain poorly understood, although several proteins, including δ-tubulin, ε-tubulin, and CPAP/SAS-4, have been implicated in these processes (Hung et al., 2000; Dutcher, 2003; Gogendeau et al., 2011; Zheng et al., 2016; Stathatos et al., 2021; Vásquez-Limeta et al., 2022; Guichard et al., 2023). Unlike the A-tubule, which appears to have a capped proximal end, the B- and C-tubules have open proximal ends, suggesting that their formation and stabilization rely on mechanisms distinct from those of the A-tubule. Interestingly, however, a previous study reported that overexpression of a γ-tubulin variant in *Tetrahymena* resulted in the loss of B- and C-tubules in basal bodies (Joachimiak et al., 2018), raising the possibility that γ-tubulin also contributes to the assembly or stability of these tubules. Consistent with this idea, we observed protofilament loss from the centriolar C-tubule in the γ-tubulin mutants. One possibility is that γ-TuRCs localized to the outer wall of the centriole contribute to the stabilization of the C-tubule, and possibly the B-tubule as well (Schweizer and Lüders, 2021; Kalbfuss and Gönczy, 2023).

In summary, we found that the *bld13* mutations in γ-tubulin lead to a partial loss of protofilaments of centriolar triplet microtubules, while the triplet structure remains stable even when some protofilaments are lost. The loss of protofilaments appears to proceed from the proximal end, potentially due to reduced minus-end capping by the γ-TuRC containing *bld13* γ-tubulin. Our findings have thus highlighted a potential role for γ-tubulin and the γ-TuRC in the stabilization of the minus end of the centriole microtubules, in addition to their established function in initiating tubulin polymerization.

## Materials and methods

### Chlamydomonas reinhardtii strains

The wild type strains CC124 and CC125, and S1-D2 were obtained from the *Chlamydomonas* Resource Center (University of Minnesota). The mutant *bld13-1* and *bld13-2* were isolated in this study by screening CC125 cells treated with N-methyl-N’-nitro-N-nitrosoguanidine (MNNG) or irradiated with UV, respectively. The isolated strains were then backcrossed twice with wild-type, and the resulting F2 progeny were used for all subsequent analyses. The diploid strains (WT/WT, WT/*bld13-1*, WT/*bld13-2*, *bld13-1*/*bld13-2*) were produced by crossing two strains with different antibiotic resistance: hygromycin and paromomycin resistance, each conferred by transformation with a plasmid pHyg3 carrying aphVII or pSI103 carrying aphVIII gene by electroporation (Ebersold, 1967; Berthold et al., 2002; Sizova et al., 2001; Yamano et al., 2013). Cells were grown in Tris-acetate-phosphate (TAP) medium with constant illumination (Gorman and Levine, 1965).

### Expression of HA-tagged γ-tubulin in *Chlamydomonas*

Genome fragments containing the γ-tubulin gene (Cre06.g299300v5, https://phytozome-next.jgi.doe.gov/) were amplified by PCR with the primers γ-tub-F (5’-AACTCCCTCACCACCCAGTCA-3’) and γ-tub-R (5’-TTTCATTAGGGACCAGGTTAG-3’) using genomic DNA of the wild-type, *bld13-1* or *bld13-2*. The forward primer (γ-tub-F) was designed to amplify approximately 500 bp upstream of the 5’UTR region of the γ-tubulin gene, thereby including its putative native promoter region within the amplified fragment. The amplified fragments were cloned into pCR-Blunt II-TOPO vector (Invitrogen). Into this plasmid, a SpeI fragment containing the aphVII gene excised from pHyg3 was inserted at the SpeI site, and a fragment containing 3×HA-tag sequences digested with NruI/ScaI from p3×HA (Silflow et al., 2001) was inserted at the site adjacent to the stop codon of the γ-tubulin gene. These plasmids were introduced into the wild-type, *bld13-1* or *bld13-2* cells by electroporation using the NEPA21 electroporator (Nepa Gene, Japan) (Yamano et al., 2013). Of the transformants that display resistance to hygromycin, the clones that express the HA-tagged γ-tubulin were screened by Western blotting using anti-HA antibody (3F10, Roche).

### Genetic mapping and whole genome sequencing

The mutations *bld13-1* and *bld13-2* were mapped to a region containing the γ-tubulin gene on Chromosome 6 by amplified-fragment-length polymorphism (AFLP)-based analyses of progenies from the mating of these mutants with S1-D2 (Kathir et al., 2003; Nakazawa et al., 2007). Recombination frequencies between each mutation and genetic markers on Chromosome 6 were determined by detecting polymorphisms in PCR products. The genetic markers used were as follows: 19-559 (5’-CAGTGTGTAGGGTTGAGCACAGGA-3’, 5’-CAGCCGGAGATGTTTACGTTTGAG-3’) 190065 (5’-GGGGCACTGACGTCAGGCAAG-3’, 5’-GCAAAGCGTCTAGCCTGGACGAA-3’, 5’-GGATACTGGACTACTTTCACGCTGCA-3’), 190071 (5’-GGCCCAGTCGATGCTGGTGTTCGG-3’, 5’-GTCTGGGTCGGCGCCCTTGAGA-3’, 5’-GGGGCACACATGATGAGAGGTGCG-3’), TUG (5’-GTCGCCAGGAATTTTGCCCCTGG-3’, 5’-GCGCGCCTGGCGGTAGCACATA-3’, 5’-AGCAGCGCTATGTTCGCTTCCCC-3’), 19-1050 (5’-AGGCAGGGGAGACATAGGTAGGAG-3’, 5’-GACCCTATGACCACCATCTGTTCC-3’), 9-0003 (5’-GTAGCACCATATGGGTGGAGTGG-3’, 5’-AGAACTGCGCCAAGATTTGCACG-3’). Of these markers, no recombinants were detected between the mutations and markers 190071, TUG, and 19-1050 in 188 mutant progenies examined.

For whole genome sequencing, genomic DNA was prepared using the DNeasy Plant Mini kit (QIAGEN) from *bld13-1* cells pretreated with gametolysin to remove the cell wall (Kindle, 1990). A total of 100 ng of genomic DNA was fragmented to an average size of ∼350 bp using a Covaris sonicator. Sequence libraries were constructed from the DNA fragments using the TruSeq Nano DNA library prep kit (Illumina), essentially following the manufacturer’s instructions. To minimize PCR amplification bias, the number of PCR cycles in the amplification enrichment step was reduced to four. The generated library was evaluated with the Bioanalyzer (Agilent) and then sequenced on a HiSeq 2500 (Illumina) in a 101 x 101 paired-end format.

The sequence reads were aligned onto the *Chlamydomonas* genome sequence version 5.3.1 (Creinhardtii_236.fa.gz) downloaded from the Joint Genome Institute (JGI) (https://phytozome.jgi.doe.gov/pz/portal) using bowtie2 (bowtie-biosourceforge.net/bowtie2/index.shtml). The *bld13-1* mutation was identified by comparing the genome sequence of *bld13-1* with that of CC124 and CC503 focusing on the mapped ∼400-kb region on Chromosome 6. The *bld13-2* mutation was identified by PCR amplification and sequencing of the γ-tubulin gene in the *bld13-2* genome.

### Western blot analysis

For immunoblotting, whole-cell extracts (10 μg/lane) from *Chlamydomonas* cells were separated by SDS-PAGE and transferred to Immobilon-P membranes (Millipore) as described in Noga et al., 2022. Anti-HA antibody bound to the membrane were detected with goat anti-rat IgG conjugated with horseradish peroxidase (HRP) (Sigma).

### Immunofluorescence microscopy

*Chlamydomonas* cells were fixed and prepared for antibody staining as described previously (Arakaki et al., 2017). Anti-HA antibody (3F10, Roche), anti-α-tubulin antibody (B-5-1-2, Sigma), Rabbit anti-SAS-6 antibody (Nakazawa et al., 2007), and anti-acetylated α-tubulin antibody (6-11B-1, Sigma) were used as primary antibodies. Goat anti-mouse IgG conjugated with Alexa Fluor 647 (Thermo Fisher Scientific), goat anti-mouse IgG(H+L) cross-adsorbed secondary antibody conjugated with Alexa Fluor 488 (Thermo Fisher Scientific), goat anti-rat IgG conjugated with Alexa Fluor 488 (Thermo Fisher Scientific), and goat anti-rabbit IgG conjugated with Alexa Fluor 546 (Thermo Fisher Scientific) were used as secondary antibodies. Images were acquired using a Nikon AX confocal microscope equipped with an NSPARC super-resolution detector (Nikon Instruments Inc.). For the observation of cytoplasmic microtubules, Z-stacks were collected at 0.15-μm intervals, and Z-projected images were generated using maximum-intensity projection of 40-50 optical sections.

### Quantification of microtubule defects

For quantification of microtubule organization (Fig. 4), Z-projected immunofluorescence images of cells stained with anti–acetylated α-tubulin antibody (Fig. 4A) or anti–α-tubulin antibody (Fig. 4B) were analyzed. Cells were categorized into four groups based on the number and organization of stained microtubules: cells with three or fewer microtubules (≤3 MTs), cells with four microtubules (4 MTs), cells with five or more microtubules (≥5 MTs), and cells with cage-like microtubules that did not originate from an obvious organizing center (CL). Statistical analysis was performed using Fisher’s exact test in GraphPad Prism to compare the distribution of microtubule categories between wild-type and mutant strains. For each strain, more than 120 cells were randomly selected and manually scored. The statistical power of 0.8 or higher was confirmed for each comparison in a post hoc manner using G*Power (Faul et al., 2007).

### Electron microscopy

Logarithmically growing cells were prefixed with 2% glutaraldehyde in 0.1 M S-collidine buffer for 1–2 hr at 0°C and postfixed with 1% OsO4 in the same buffer for 1-2 hr (Wright et al., 1983). The specimens were then dehydrated and embedded in EPON 812. Thin sections (50–70 nm) were stained with aqueous uranyl acetate and Reynold’s lead citrate and observed with JEM-1400plus at 80 kV or JEM-1230R at 100 kV (JEOL).

### Analysis of protofilament loss in centriolar triplet microtubules

To evaluate triplet microtubule integrity, we collected electron microscope images of triplet microtubules in the distal, central and proximal regions of the A tubule, and the central region of the C-tubule of *bld13-1* and *bld13-2* centrioles (Table 2) based on the clarity of the protofilaments. Potential intrinsic variation among the nine triplets within each centriole was not considered during image selection. To evaluate protofilament loss, each tubule image was divided into sectors (12 for the A-tubule and 6 for the C-tubule), and the number of missing protofilaments in each sector was scored (Fig. 5C). Missing protofilaments were scored in sectors rather than in numbered protofilaments because the images often lacked sufficient resolution to identify specific protofilament numbers. The B-tubule did not show any protofilament loss.

To assess whether the distribution of protofilament loss across sectors was uniform or showed a preferential orientation, circular statistics were employed. Specifically, the Hodges-Ajne test was performed for each genotype to test for uniformity on the circular data using a custom MATLAB script. Sample sizes were initially determined based on experimental feasibility. The statistical power of 0.8 or higher was confirmed for each comparison in a post-hoc manner using G*Power (Faul et al., 2007).

## Acknowledgements

We thank Dr. Ritsu Kamiya (Chuo University) for critically reading the manuscript, Dr. Takako Kato-Minoura (Chuo Univ.) and OIST Imaging Section for use of electron microscopes, OIST Memory Research Unit, OIST Evolution, Cell biology, Symbiosis Unit, OIST Membrenology Unit, Dr. Midori Ohta (OIST), Dr. Petra Svetlikova (OIST), Dr. Yohsuke Moriyama (OIST) and Dr. Akane Furuta (NICT) for helpful advice and fruitful discussions. This work was supported by JSPS KAKENHI (16J40154 to YN, 23K23905 to KW, and 21K06257 and 24K02055 to MH). This work was also supported in part by NIBB Collaborative Research Program (14733) to KW.

## Author contributions

Conceptualization: Yuki Nakazawa, Mao Horii, Masafumi Hirono

Methodology: Yuki Nakazawa, Mao Horii, Masafumi Hirono

Investigation: Yuki Nakazawa, Naoto Kubota, Mao Horii, Akira Noga, Yoshikazu Koike, Katsushi Yamaguchi, Shuji Shigenobu

Validation: Yuki Nakazawa

Formal analysis: Yuki Nakazawa, Hideo Dohra

Resources: Yuki Nakazawa, Hideo Dohra, Katsushi Yamaguchi, Shuji Shigenobu, Ken-ichi Wakabayashi, Masafumi Hirono

Data curation: Yuki Nakazawa, Hiroko Kawai-Toyooka, Masafumi Hirono

Writing – original draft: Yuki Nakazawa, Masafumi Hirono

Writing – review & editing: Yuki Nakazawa, Masafumi Hirono, Ken-ichi Wakabayashi

Visualization: Yuki Nakazawa, Hiroko Kawai-Toyooka

Supervision: Masafumi Hirono

Project administration: Yuki Nakazawa, Masafumi Hirono

Funding acquisition: Yuki Nakazawa, Ken-ichi Wakabayashi, Masafumi Hirono

## Data availability

The datasets supporting the findings of this study will be deposited in Dryad upon acceptance and will be made publicly available prior to publication.

---

**Fig. S1.**
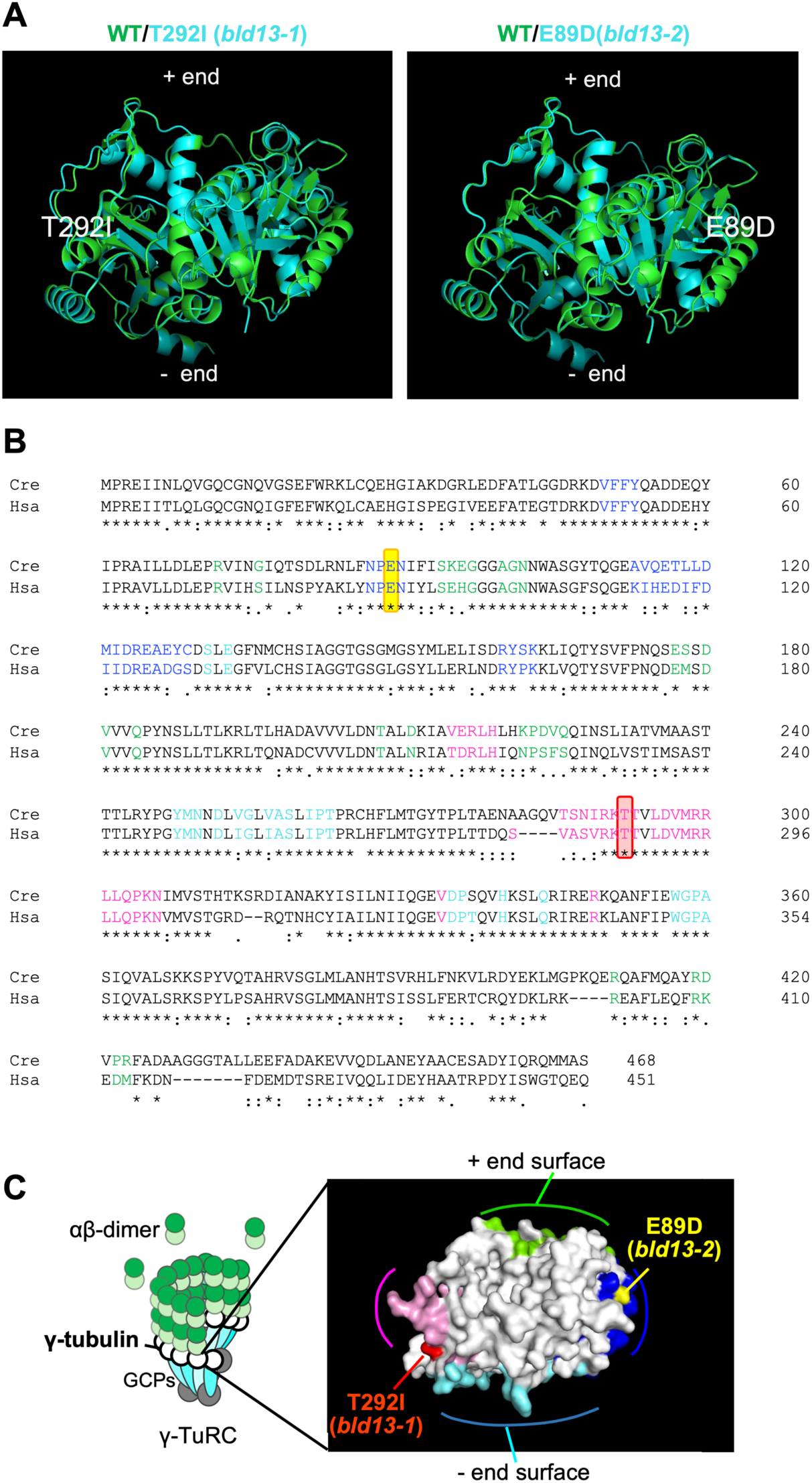
Effects of the *bld13* mutations on the γ-tubulin structure predicted by AlphaFold2. (A) AlphaFold2 models of the *bld13-1* and *bld13-2* γ-tubulin molecules (blue) are merged with that of the wild-type (green). The mutation sites (T292I and E89D) are shown in white. (B) An amino acid sequence alignment of human γ-tubulin (Acc. No. NP_001061.2) and *Chlamydomonas* γ-tubulin (Acc. No. AAA82610.1). Amino acids homologous or conserved between the two proteins were identified by Clustal Omega (Sievers and Higgins, 2014) and indicated by an asterisk or dot, respectively. Indicated by color are the amino acids located on the + end surface (green) and the - end surface (light blue), and those close to the adjacent γ-tubulin molecules (blue and pink). The positions of the mutated amino acids (E86 and T292) are indicated with yellow and red boxes. Note that they are one of the amino acids located close to the adjacent γ-tubulin. (C) Positions of the amino acid changes on the γ-tubulin 3-D structure. A model of the human γ-tubulin 3-D structure, based on the PDB file 1Z5W (Aldaz et al., 2005), is generated using PyMOL (right). This model shows the surface of the γ-tubulin molecule in the same orientation as in the γ-TuRC (left). The colors correspond to those in (B).

